# Hierarchical Evolutionary Preferences Explain Discrepancies in Expected Utility Theory

**DOI:** 10.1101/081570

**Authors:** Michael Holton Price, James Holland Jones

**Affiliations:** Santa Fe Institute; Department of Earth System Science, Stanford University; Woods Institute for the Environment, Stanford University; Department of Life Sciences, Imperial College London

## Abstract

The standard axiomatic theory of rationality posits that agents order preferences according to the average utilities associated with different choices. Expected Utility Theory has repeatedly failed as a predictive theory of choice behavior, as reflected in a growing literature in behavioral economics. Evolutionary theorists have suggested that seemingly irrational behaviors in contemporary contexts may have once served important functions, but there has been little attempt to formalize the relationship between evolutionary fitness and choice behavior. Biological agents should optimize fitness, but fitness itself is not a reasonable value function for decision-making since its time-scale exceeds the lifespan of the decision-maker. Consequently, organisms use proximate motivational systems that work on appropriate time-scales and are amenable to feedback and learning. We develop an evolutionary principal-agent model in which individuals maximize a set of proximal choice variables (age-specific demographic rates), the interests of which are aligned with fitness. The solution to our model yields pessimistic probability weightings compatible with the Rank-Dependent Expected Utility family of choice models. The pessimistic probability weighting characteristic of these models emerges naturally in an evolutionary framework because of extreme intolerance to zeros in multiplicative growth processes.

## Introduction

A herder in in the arid north of Kenya has to make decisions about the composition of his herd [1, 2, 3]. Should he restrict his herd to small, high-productivity goats or, when he gets enough goats, should he trade them for heartier, slower-breeding camels? The growth rate of goat herds, even accounting for their increased susceptibility to high mortality during drought years, is three times that of camels and the expected energetic payoff to keeping goats – more precisely, the instantaneous rate at which wealth accumulates – is therefore substantially higher than a mixed herd of goats and camels. However, pastoralists in the region favor mixed herds and will even increase the fraction of camels dramatically above a critical wealth threshold [1, 2]. Are these herders engaging in irrational economic decision-making by under-valuing the payoff of an all-goat herd? While the expected return is lower for a mixed herd of predominantly camels, households pursuing this strategy have a much higher probability of long-term persistence, nearly 70%, compared to a 0.4% probability of persistence for the goats-only strategy. Why is this so?

In part, mixed herding diversifies risk. More importantly, however, the growth of these herds, like any biological growth process, is mutliplicative and the rate of increase is stochastic. Furthermore, the herder faces more than just a decision about long-term wealth accumulation (after all, allocation of investment when growth is stochastic and multiplicative is, a well studied topic in finance [e.g., 4] and economics [e.g., 5], so we wouldn’t have much to say). In particular, herd products and income from the herd provide each household’s primary yearly consumption, and consequently there are two processes that matter to the herder: herd growth and household consumption. We might better interpret the herder’s decision as not about herd management, *per se*, but rather about household consumption through time, for which herd dynamics can lead to very uneven yearly consumption.

Like growth of the herd, household survival can be thought of as a stochastic gamble, in which the survival probability is a function of household consumption. For such a multiplicative process, a severe shortfall in any time period that leads to a low survival probability can never be compensated for by high survival in other time periods (even a large number of other time periods). A virtually identical perspective arises when considering the evolutionary fitness of individuals because, as we will describe and formalize, fitness is determined by stochastic, multiplicative processes; most obviously, individual survival through time is a component of fitness, and like household survival, individual survival is a binary outcome variable with differing probabilities each time period. In this article, we explore what this means for the evolution of organism’s decision making.

Organisms, humans included, must make decisions about foraging, reproductive, social, and political behaviors that have consequences for proximate outcomes such as satiety, income, wealth, happiness, sexual satisfaction, and well-being. These decisions also have consequences for ultimate outcomes such as fitness and, consequently, there are strong expectations that selection acting on differential fitness will shape the decision-making system. The rational-choice tradition suggests that individuals make decisions that maximize some objective function typically denoted by the catch-all term “utility” [6]. In the presence of variability in outcomes, the objective function is the expectation of utility. In particular, a set of possible payoffs (i.e., a “lottery”) **x** = [*x*_1_, *x*_2_, …, *x*_*n*_]^*T*^ with associated probabilities **p** = [*p*_1_, *p*_2_, …, *p*_*n*_]^*T*^ is preferred to some other lottery **y** = [*y*_1_, *y*_2_, …, *y*_*n*_]^*T*^ with probabilities **q** = [*q*_1_, *q*_2_, …, *q*_*n*_]^*T*^ if some value function that associates payoffs with probabilities *V* (**x**; **p**) is greater than that associated with *V* (**y**; **q**): *V* (**x**; **p**) > *V* (**y**; **q**). In the absence of other information, a reasonable value function to associate payoffs with probabilities is linear in the probabilities and utilities of distinct outcomes, e.g., *V* (**x**; *P*) = Σ_*i*_*p*_*i*_ *u*(*x*_*i*_). The decision model in which preferences are ordered by their expected utilities is known as expected utility theory (EUT) and was first formalized in its modern form by von Neumann and Morgenstern [6].

The EUT approach has been very fruitful for economics, political science, and other behavioral sciences. Like many economic theories it is axiomatic and the fundamental axioms that underlie EUT (completeness, transitivity, continuity, and independence) are sensible requirements that ensure that preferences are consistent [6, 7]. However, an enormous literature has developed showing that people violate the axioms underlying EUT in both experimental and naturalistic contexts. Some examples include the common consequence effect (Allais paradox), the common ratio effect [8], ambiguity aversion [9], preference reversals between gambles represented as bids versus choices [10], the incommensurability of risk-sensitive behavior for high-vs. low-stakes gambles [11], and abundant evidence that framing and reference points induce departures from canonically-predicted behavior [12, 13, 14].^1^

Machina [16, 17] noted there is nothing inevitable about expected utility serving as the objective function for preference ordering and suggested that nonlinear functions mapping values and utilities might account for these types of systematic departures from the predictions of EUT. What is less clear is what mechanistic or functional basis such nonlinear functions might have. One possibility for the origin of such nonlinear functions is evolutionary history. While many economists and decision theorists express disdain (or at least indifference) to the possible evolutionary origins of decision biases, there is a thread of evolutionary logic that runs through parts of the behavioral-economic literature. For example, an interpretation of Kahneman and Tversky’s dual-process model, as articulated by Kahneman [18, p. 697], suggests that “intuitive judgments occupy a position – perhaps corresponding to evolutionary history – between the automatic operations of perception and the deliberate operations of reasoning.” Various well-known decision biases – particularly loss-aversion – have been interpreted in light of evolutionary success [19, 20]. The nature of risk determines the degree to which EU maximization is evolutionarily optimal. As Robson [21] notes, evolutionarily-derived preferences over commondities can lead to a situation where a gamble involving idiosyncratic risk is preferred to one with aggregate uncertainty, even though the former is stochastically dominated by the latter from an individual standpoint. Netzer [22] suggests that selection favored distinct utility functions of short-run and long-run decisions, leading to intertemporal inconsistencies in time preferences. The putatively superior performance of some decision biases under ecological conditions has been argued for by various authors [23, 24]. Thus, there is a thread of scholarship suggesting that behaviors that appear irrational in contemporary contexts may have served important functions during human evolutionary history, but there has been little attempt to formalize the relationship between evolutionary fitness and choice behavior. In line with Machina’s observation, we describe and interpret a formal model of the evolution of preferences that leads to decision-making rules which violate EUT but nevertheless maximize organisms’ evolutionary fitness. In particular, organisms display a profound intolerance for zeros or low values in key evolutionary parameters. This intolerance for zeros arises from a fundamental difference between economic utility and fitness. Cumulative expected utility is additive across time periods, whereas fitness is multiplicative [25, 26]. Within an individual’s lifetime, the individual must survive each previous time period to reach a given age to reproduce. Across generations, individual lineages must persist; referring back to the opening example of herd management, household persistence is crucial to lineage persistence. Zeros in such processes represent absorbing states. Consequently, risky strategies that may be acceptable under an additive metric such as cumulative expected utility can be unacceptable, and even catastrophic, given a multiplicative metric such as survival or lineage persistence.

We present a model of the evolution of decision making in which organisms can adopt EUT-type risk preferences toward lotteries, then show how the type of evolutionary conservatism described in the preceding paragraph leads to pessimistic subjective probability weights when rank-dependent expected utility theory (RDEUT) replaces EUT in the evolutionary model. We focus on RDEUT for two reasons. First, it is an appealing and popular theoretical generalization of EUT. Machina [27, p. 1237], for example, describes RDEUT as “the most natural and useful modification of the classical expected utility formula.” RDEUT allows the nonlinear probability weighting that is commonly observed in choice experiments but avoids the problem of stochastic dominance in classical Prospect Theory [12]. Second, RDEUT is one of the core components of cumulative prospect theory (CPT), along with loss aversion [13, 28]. Consequently, RDEUT, either in its own right or as a component of CPT, is one of the most important contemporary generalizations of EUT [29, 28]. Although we focus on RDEUT in our model, we consider our conclusions to have greater scope since subjective probability weighting is a general topic in the literature on the psychology of risk [30].

## Natural Selection and Preferences

We are interested in economic decisions in the broadest sense. At the basic level, organisms are making decisions over what can be thought of as different lotteries. For example, should a herder focus on one animal or create a mixed herd? Should a forager hunt for sand monitor lizards or hill kangaroo [31]? Should a peasant farmer intensify cultivation of a nearby garden plot or spread effort across two geographically distinct plots [32]? Should a woman wean her infant and have another baby or continue nursing and delay reproduction [33, 34]? Should an individual buy or rent a home [35]. In these examples, each lottery yields different payoffs probabilistically. All of these examples have a clear impact on fitness, but it is extremely unlikely that any conscious fitness-maximization goal plays significantly into any of the decision-makers’ choices. Instead, their decisions are shaped by preferences over a variety of proximate currencies like hunger/satiety, feelings of security, or feelings of love and responsibility for children. Samuelson and Swinkles [36] raise the important question, given the evolutionary mandate to successfully leave descendants, why do people have preferences for anything but fitness?

### Why Have Preferences for Proximate Quantities?

In the substantial literature that addresses the question of why people have preferences for proximate currencies rather than fitness itself [37, 38, 39, 40], three inter-related factors loom largest. First, natural selection operates on time-scales that are longer than the lifespans of the organisms whose behavior it shapes and, furthermore, selection is a stochastic, undirected process. Second, organisms regularly encounter novel situations for which natural selection is unable to directly specify behaviors. Third, the types of solutions that might emerge via natural selection to address the first two factors are constrained by trade-offs imposed by the cost of gathering and processing information. These observations can be accommodated within a single analytic framework by utilizing the economic concept of the principal-agent problem [38] which will allow us to address the question of why organisms have preferences defined over proximate currencies, rather than the ultimate currency of fitness.

### The Evolutionary Principal-Agent Problem

Consider a principal that possesses certain goals that it is unable to achieve unless it acts through agents over which it has only indirect control [38]. Natural selection is the ultimate arbiter of which biological entities remain and increase in a population. While the process of natural selection lacks agency, it is nonetheless useful to consider the outcomes of selection as having been designed [41]. It is in this sense that natural selection can be thought of as a principal with a goal of maximization of fitness. However, there are clear limitations in the ability of selection to achieve a solution, as enumerated above. Due to the obvious lack of direct control, selection shapes cognitive mechanisms which are, on the whole, consistent with fitness maximization. As noted by Binmore [38], these collective proximate cognitive mechanisms can be thought of as the agent in the evolutionary principal-agent problem, wherein the principal (selection) “seeks to design an incentive scheme that minimizes the distortions resulting from having to delegate to the agents” [38]. The extent to which selection minimizes these distortions for a given organism in a given setting depends on the structure and strengths of the constraints it faces.^2^

Figure 1 encapsulates the conceptual model that emerges from the principal-agent framework. At the bottom level, an individual is choosing between lotteries that influence the outcomes **x**_**1**_, **x**_**2**_, …. The outcomes of these lotteries contribute to utilities, *u*_*j*_, at the next level. These lotteries can be thought of either as motivational systems such as satiety, sexual gratification, or happiness (akin to the classical notion of utility) or as proximate determinants of fitness such as infant survival or total fertility. What links these two different notions of “utility” is that they work on a time-scale where the organism can use feedback from outcomes to change its preferences and therefore decision-making. These proximal utilities then contribute ultimately to fitness.

**Figure 1:**
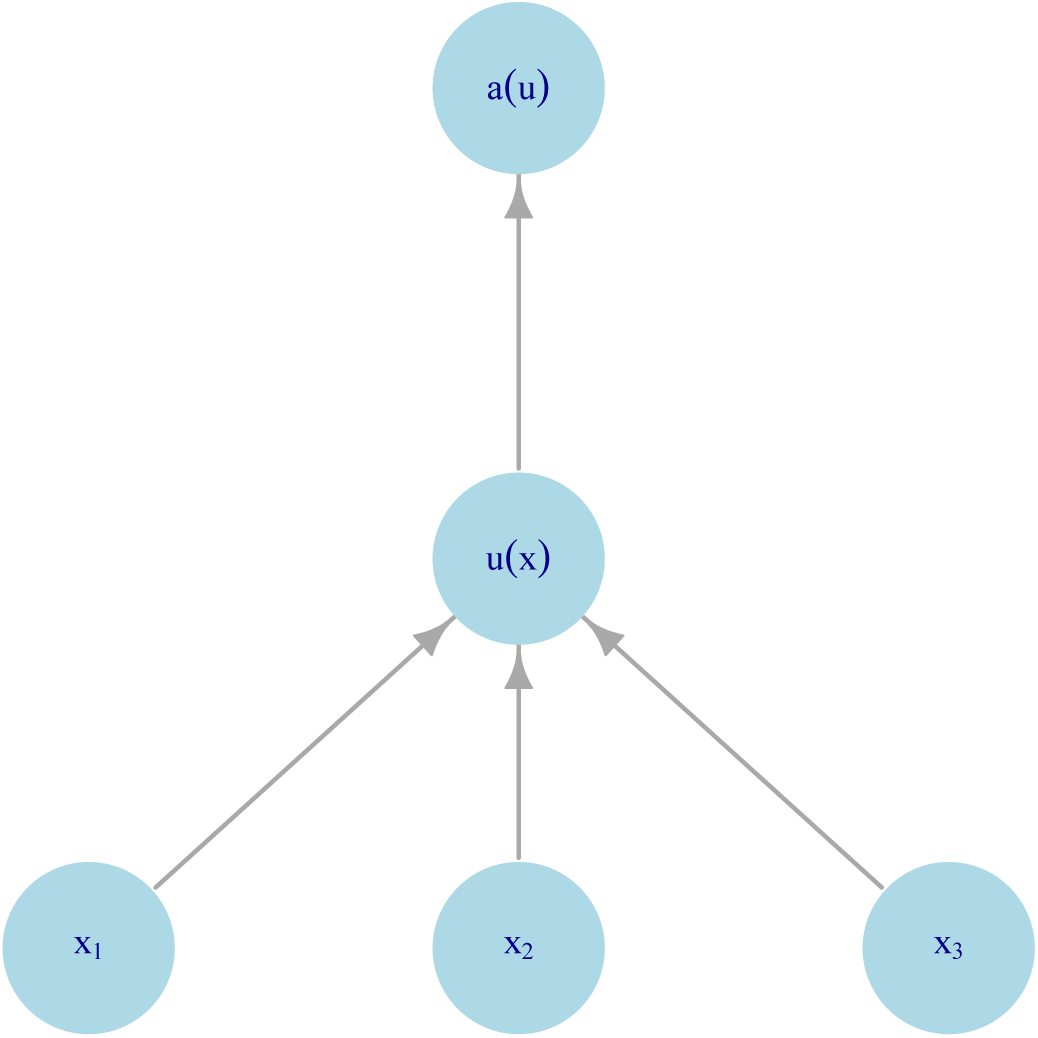
Hierarchical model of decision-making as a principal-agent problem. Input variables (**x**_**i**_) contribute to intermediate utility (*u*(*x*)), which themselves contribute to fitness (*a*(*u*)).

### Applying the Hierarchical Framework

The preceding section describes a conceptual framework for understanding the evolution of economic preferences. The main conclusion is that the Evolutionary Principal-Agent Problem implies the existence of proximate decision making mechanisms within a hierarchical framework. We anticipate that this will be a relatively uncontroversial perspective. We anticipate greater controversy in the particular predictions of specific models that implement the hierarchical framework.

In this section, we describe how to implement a formal model in accordance with this conceptual framework. In so doing, we provide a high-level summary of the actual model described below upon which our results and conclusions are based.

Three elements are needed to apply the conceptual framework encapsulated by Figure 1: (1) an evolutionary model that defines how fitness depends on some determinants of fitness and how those determinants of fitness depend on consumption allocated to them; (2) an economic model of decision making; and (3) a mechanism to link the evolutionary and economic models and thus provide novel predictions or insight into behavior.

#### Evolutionary model

The evolutionary formalism we use is stochastic age-structured life history theory as described by Tuljapurkar [45]. We assume a randomly mating, diploid population. Different phenotypes make different consumption decisions when faced with risk and environmental uncertainty. These differing consumption streams lead to different realizations of age specific survival and fertility and thus to different growth profiles for the phenotypes. Tuljapurkar [45] shows that the logarithmic growth rate of a phenotype governs invade-ability (formalized below). In terms of Figure 1, the logarithmic growth rate is the measure of fitness whereas age specific survival and fertility – which change through time – are the determinants of fitness.

#### Economic model

The economic formalism we use is rank dependent expected utility theory (RDEUT). RDEUT is very similar in form to Expected Utility Theory. In EUT, the value of a lottery is

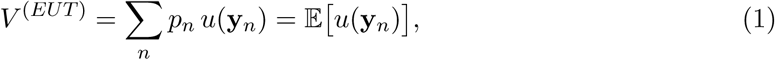

where *p*_*n*_ is the probability of an outcome with payoff *y*_*n*_ and *u*(*y*) is a utility function that depends only on the payoff amount. In RDEUT, the value of a lottery is

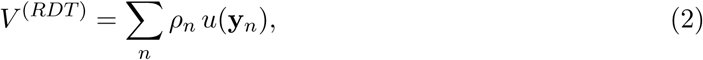

where *ρ*_*n*_ is a re-weighting of the probability vector *p*_*n*_. At this point, three properties of this re-weighting should be emphasized. First, it is implicitly assumed that the outcomes are ordered from most to least desirable (*y*_1_ > *y*_2_ > *y*_3_ …). Second, ***ρ*** is normalized to sum to 1 just like a true probability vector, 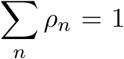. Third, although ***ρ*** depends on the ordering of outcomes it does not depend on the actual values of those outcomes aside from this ordering, ***ρ*** = ***ρ***(**p**) where the aforementioned implicit ordering of *y*_*n*_ applies.

The final high-level aspect of RDEUT that is useful to know at this stage of explication is the concept of pessimism/optimism. In RDEUT, a pessimistic decision maker utilizes a probability re-weighting that ascribes a lower value than the EUT valuation to any gamble: *V* ^(*RDT*)^ < *V*^(*EUT*)^. An optimistic decision maker ascribes a higher value: *V* ^(*RDT*)^ > *V* ^(*EUT*)^. RDEUT also allows for no re-weighting, *V* ^(*RDT*)^ = *V* ^(*EUT*)^ and equivalently *ρ*_*n*_ = *p*_*n*_. Thus, RDEUT generalizes EUT and recovers EUT in the appropriate limiting case.

#### Linking the models

To link the models, we assume that the determinants of fitness in the evolutionary model (age-specific survival and fertility) can be associated with the utility function in EUT. That is, at least to first order, an organism makes proximate decisions based on the expected values of the determinants of fitness. Other assumptions can be made to link the models, but we consider this an eminently sensible assumption to make given the limitations implied by the Evolutionary Principal-Agent Problem. It seems much easier for an organism to reason about the mean number of offspring a given strategy will yield (perhaps with a time discount factor applied on the timing of reproduction) than for an organism to reason about the ultimate fitness consequences of a given strategy.

However, we will show that averaging survival or fertility is systematically “wrong” – i.e., leads to lower fitness – when stochastic uncertainty in achieved outcomes is accounted for. Specifically, the true fitness of a strategy that involves stochastic uncertainty is lower than that implied by the corresponding mean strategy that accounts for only mean survival and fertility. This is precisely the condition for pessimism in RDEUT, and implies that an organism can “correct” its expected utility bias by re-weighting the probabilities of a gamble as in RDEUT. In addition to establishing a general result – that there is always a level of pessimism that is superior to no probability re-weighting (Section) – we present a two specific models in which the level of pessimism (or optimism) evolves, and demonstrate that the highest fitness is achieved by an intermediate level of pessimism (Sections and).

## A Model of the Evolution of Pessimism

To model the population dynamics and genetics, we utilize the formalism described by Tuljapurkar [45]. We assume a diploid locus with alleles 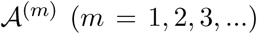, random mating, a fixed sex-ratio, and ignore sex-differences. The phenotypes 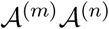 represent distinct behavioral strategies that interact with the environment to determine an individual’s life-history traits in each time period. Tuljapurkar [45] derives two essential results. First, he shows that the logarithmic growth rate of phenotype 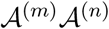 governs invade-ability. That is, a population of homozygotes with allele 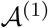 is resistant to invasion by allele 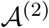 if *a*^(11)^ *> a*^(12)^, where *a*^(*mn*)^ is the logarithmic growth rate of phenotype 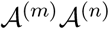. Second, Tuljapurkar provides an analytic formula for the logarithmic growth rate *a* when the variance in life history traits is small. The fundamental tool on which these results rely is the Leslie matrix; we introduce the Leslie matrix in the in the next section, then (a) describe how stochastic uncertainty can be added, (b) introduce the economic theory, and (c) link the evolutionary and economic theory to establish our central result, which is that natural selection will favor behavioral phenotypes that utilize pessimistic subjective probability weights to value gambles.

### Life history Traits

Consider an age structured population where each age class *j* accounts for individuals between *y*_*j*_ = *j* ∆*y* and *y*_*j*+1_ = (*j* + 1) ∆*y* years of age. The key demographic traits that govern the population’s dynamics are the age-specific fertility, *F*_*j*_, and the age-specific survival probabilities, *P*_*j*_. *F*_*j*_ is the mean number of offspring that an individual in age class *j* contributes to the juvenile age class (*j* = 1) going from time *t*_*n*_ to time *t*_*n*+1_ = *t*_*n*_ + ∆*y*, where *n* indexes time steps. *P*_*j*_ is the probability that an individual will survive from age class *j* to age class *j* + 1. The Leslie matrix, **A**, has the age-specific fertilities on the first row and the age-specific survivals on the sub-diagonal,

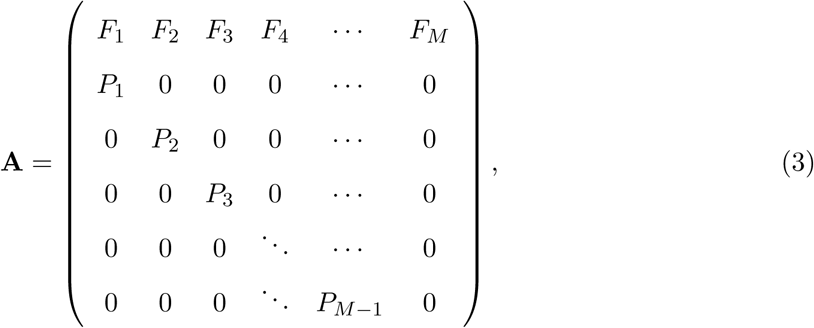

where *M* is the number of age classes. The Leslie matrix projects the population vector **z**_*n*_ one time step into the future, **z**_*n*+1_ = **Az**_*n*_. We assume that the Leslie matrix is irreducible and primitive, so that **A**^*n*^ converges to a stable age distribution for large enough *n*. Given these assumptions, there exists a unique dominant eigenvalue *λ* that is the per-period growth factor of the population. In the absence of stochastic fluctuations of the Leslie matrix elements, *a* = log *λ* (stochasticity is considered below). Hence, a population of homozygotes with phenotype *A*^(1)^*A*^(1)^ is resistant to invasion by allele *A*^(2)^ if *λ*^(11)^ *> λ*^(12)^.

Associated with *λ* are the dominant left and right eigenvectors of **A**, which we represent by ***ν*** and ***ω***, respectively. ***ν*** is the vector of age specific reproductive values and ***ω*** is the stable age distribution vector.

### Stochastic Uncertainty

We now generalize the preceding material to allow for stochastic uncertainty in the Leslie matrix elements. The crucial idea is that the Leslie matrix is not fixed from one time step to another, but instead depends on the changing state of the world. Following Tuljapurkar [45], consider an environmental sequence *ε* with an associated sequence of Leslie matrices **A**_1_, **A**_2_, …, **A**_*n*_. Let

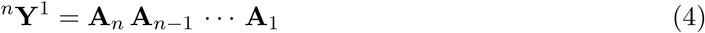

represent the product matrix which governs population growth to time period *n*, and let *λ*_*n*_, ***ω***_*n*_, and ***ν***_*n*_ represent, respectively, the dominant eigenvalue, corresponding right eigenvector, and corresponding left eigenvector of ^*n*^**Y**^1^. Let *λ*, ***ω***, and ***ν*** represent the corresponding quantities for the mean Leslie matrix. In mathematical jargon, this is abuse of notation since ^1^**Y**^1^ is not typically the same as the mean Leslie matrix and ***ω***, etc., have already been defined with respect to the non-stochastic Leslie matrix. However, the meaning of symbols is always clear from context. Without loss of generality, we assume that ***ω***_*n*_ and ***ν***_*n*_ are normalized to 1. Weak ergodicity guarantees certain results. First, it guarantees that

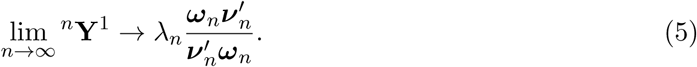

Second, it guarantees the convergence of age structure to a stable age structure for any non-negative and non-zero starting vector **z**_0_. To formalize this, let

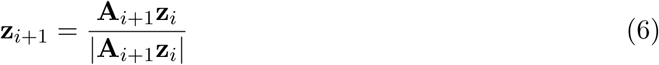

be the normalized age structure at time period *n*. Then

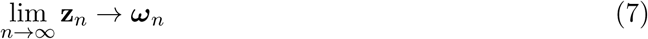

An analogous result holds for 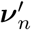 with left multiplication. The logarithmic growth rate is

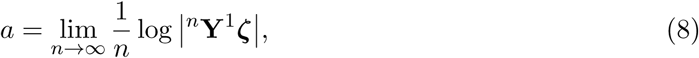

where ***ζ*** is an arbitrary initial age structure. Stochasticity lowers the logarithmic growth compared to a non-stochastic reference Leslie matrix with the same mean life history traits. Effectively, this is because growth is a multiplicative process and the geometric mean is always less then the arithmetic mean [25]. Tuljapurkar [45] derives a useful “small noise approximation” for the logarithmic growth rate assuming small fluctuations of the Leslie matrix elements relative to the mean values,

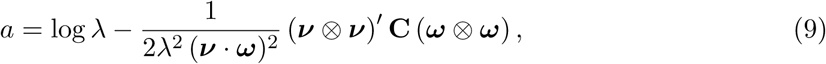

where **C** is the covariance matrix of **A**, *⊗* is the Kronecker product, and the serial autocorrelation term in Tuljapurkar’s approximation is not included. If only one Leslie matrix element is considered, Equation 9 simplifies to

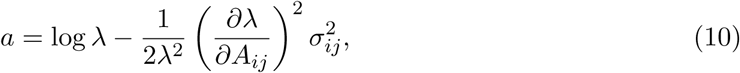

where 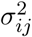 is the stochastic variance associated with matrix element *A*_*ij*_. Equation 10 is the key evolutionary result. In the next section, we discuss the relevant economic theory.

### Rank-Dependent Expected Utility Theory

Expected utility, combined with exponential time discounting of delayed outcomes, represents the conventional economic model of decision making. Expected Utility Theory follows from four axioms that ensure consistent preferences over risky prospects: completeness, transitivity, continuity, and independence. However, as noted in the Introduction, there are numerous mismatches between the expectations of EUT and observed choice behavior. Many of these empirical violations involve the independence axiom. It was therefore seen as desirable to recast a theory of choice that kept the desirable properties of EUT, such as the transitivity of preferences and the ability to combine bundles of goods as afforded by continuity, but did not rely on the difficult-to-justify requirement that decisions be independent of (possibly) irrelevant information.

A notable example of the violation of the independence axiom – and perhaps even the most notable empirical violation of EUT – is the Allais paradox, or common consequence effect. The independence axiom asserts that adding a common outcome to two prospects should not change the preference ordering of the prospects. Quiggin [46] writes: “The Allais problem is the *pons asinorum* of theories of choice under uncertainty. Almost all of the many authors who have introduced new models of choice under uncertainty in the last ten years [i.e., alternatives to expected utility theory] have included a demonstration that the model is consistent with the behavior revealed in this problem.”

Subjective probability weighting has been suggested as one way to accommodate the Allais paradox. In the simplest formulation of probability weighting^3^, the weights which enter the cumulative utility sum for probabilistic outcomes are a direct function of the probability of each individual outcome,

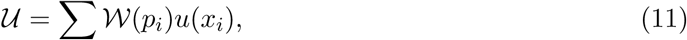

where 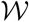 is the weighting function and *p*_*i*_ is the probability of outcome *i*. 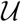 is used instead of *U* in Equation 11 so *U* can be reserved for expected utility sums (which Equation 11 is not, despite its similar form). This straightforward method of probability weighting was adopted in the original formulation of Prospect Theory (PT [12]), one of the first generalizations of EUT to gain widespread attention. However, it was recognized that PT violated first order stochastic dominance, which occurs if one gamble is preferred to another, even though the second gamble is strictly better than the first in the sense that the second gamble’s cumulative density function is at least equal to that of the first over its entire domain, and is strictly greater over at least part of the domain. An ad hoc fix of the PT violation of stochastic dominance was incorporated via an editing process that occurred before the probability weighting occurred. In addition to lacking parsimony, this editing process can induce intransitivity in pairwise choices [48]. The appeal of PT was its selective incorporation of the more desirable features of EUT, while nevertheless rejecting the independence axiom, so that the Allais paradox and other behavioral violations could be accommodated, but a model that leads to the favoring of stochastically-dominated choices or intransitivity was clearly unacceptable.

Following the formulation of PT, a number of scholars began to formulate theories that adopted the three less controversial axioms of EUT (completeness, transitivity, and continuity) while simultaneously avoiding the violation of stochastic dominance exhibited by PT in its unedited form. The culmination of this work was the simultaneous and independent discovery of RDEUT by at least three scholars [48, 49, 50]. RDEUT is the only axiomatic formulation of utility theory that simultaneously satisfies completeness, transitivity, and continuity while avoiding the violation of stochastic dominance. Violations of stochastic dominance can only be avoided if the subjective probability weighting depends in a quite specific manner on the entire vector of probabilities for a prospect. Assume that the probabilities of some prospect are sorted from most desirable to least desirable according to the values of their corresponding outcomes (*x*_1_ ≥ *x*_2_ ≥ *x*_3_ ≥ …). Next, define the cumulative probability, or rank [28], as

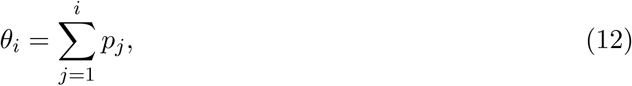

where *θ*_0_ = 0. *θ*_*i*_ is the probability of receiving an outcome that is as good or better than *i*. The probability weighting on outcome *i* is given by

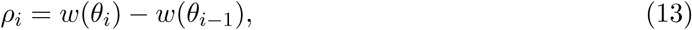

where *w* is the weighting function defined on ranks. *w* is used rather than 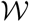 to emphasize that *w* is defined on probability ranks, whereas 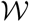 is defined on probabilities. *w* maps the interval [0, 1] onto itself and must be an increasing function of *θ*. The rank dependent expected utility is

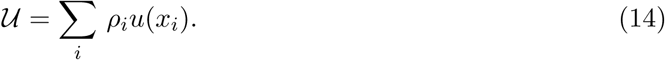

Figure 2 plots three different probability weighting functions. The middle curve is a standard form that exhibits two suggested biases in the psychophysics of probability distortions: likelihood sensitivity and pessimism [30, 28]. Likelihood sensitivity occurs because differences near the endpoints of the probability scale (0 and 1) loom larger than differences in the interior of the interval. Conversely, differences in the middle of the scale loom smaller, so that the difference between 0% and 1% is psychologically much more salient than the difference between 60% and 61%. To account for diminished likelihood sensitivity for intermediate probabilities, the probability weighting function must have a inverse-S shape so that the center is more flat than the endpoints.

**Figure 2:**
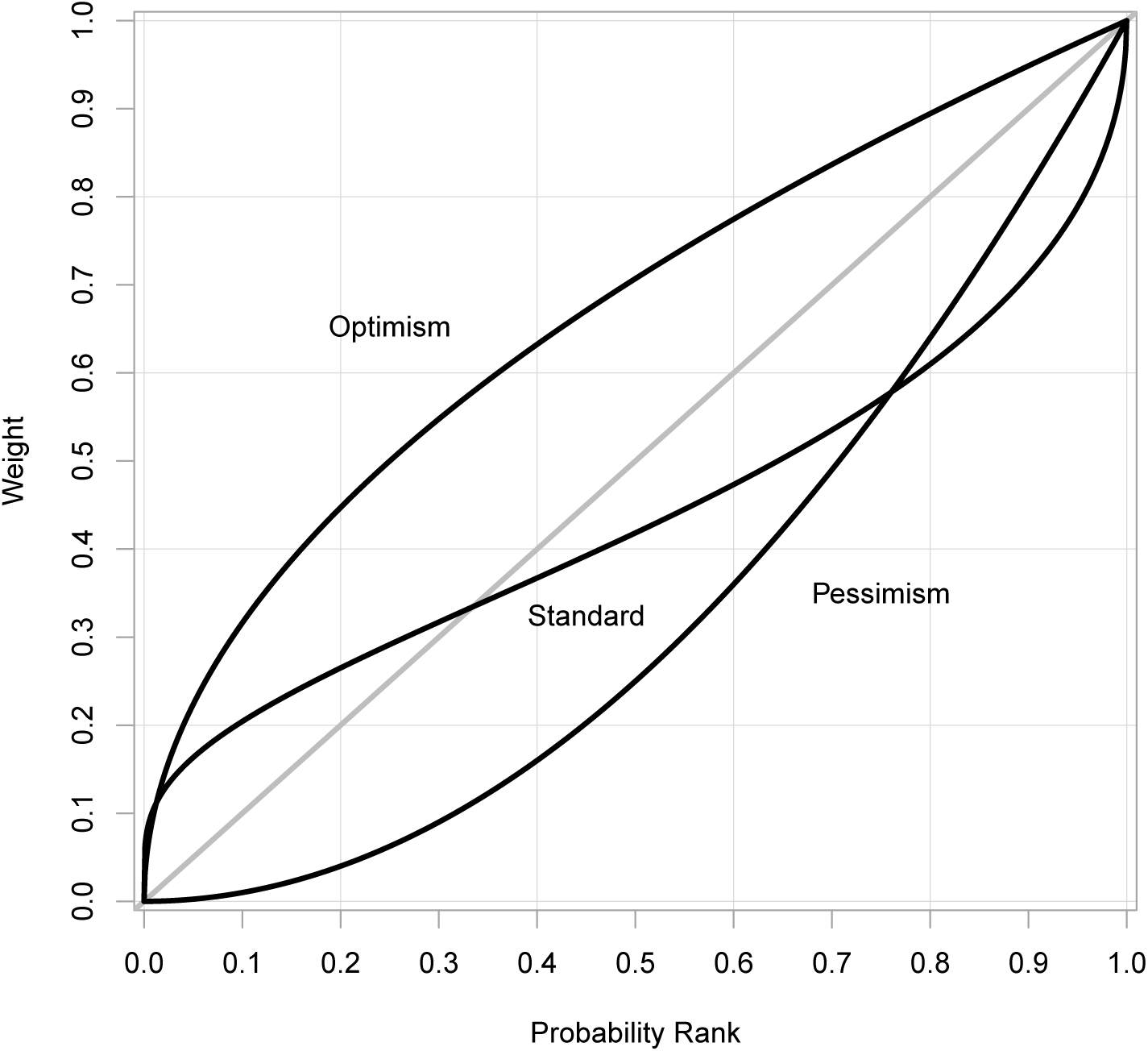
Probability weighting functions for the RDEUT model. The top curve exhibits optimism over its entire domain. The bottom curve exhibits pessimism. The middle curve is a standard curve adopted in much of the recent literature; it combines pessimism with likelihood sensitivity.

Pessimism involves the under-weighting of desirable outcomes relative to their probability of occurrence. Two types of pessimism can be defined relative to the probability weighting function, *w*(*θ*): regular pessimism and strong pessimism [46]. Regular pessimism (or simply pessimism) occurs where *w* is below the identity line *w* = *θ*, whereas optimism occurs where *w* is above the identity line. Strong pessimism occurs where *w* is convex (i.e., *w*″ > 0), whereas strong optimism occurs where *w* is concave (i.e., *w*″ < 0). For many functional forms of *w*, regular pessimism and strong pessimism occur on the same or nearly the same intervals (and similarly for optimism).

RDEUT accounts for the Allais paradox through pessimistic probability weighting. Segal [51], for example, has formulated a general statement of the Allais paradox using arbitrary payoffs and non-discrete probability distributions, and shown that RDEUT with a probability weighting function *w* exhibiting strong pessimism on its entire domain (i.e., the unit interval) can accommodate any and all formulations of the generalized Allais paradox. However, only a subset of the generalized formulations are encountered in daily life or economic choice experiments, so less restrictive probability weighting functions can accommodate the empirically relevant cases, although some pessimism is still a necessary element. For example, as already discussed, pessimism is one of the two primary components of the standard probability weighting curve (the other being likelihood sensitivity), which exhibits pessimism over only part of its domain. Consequently, we utilize regular pessimism as opposed to strong pessimism as the relevant definition of pessimism.

When it was realized that RDEUT avoided violations of stochastic dominance and could accommodate the Allais paradox, PT was reformulated to include RDEUT, becoming Cumulative Prospect Theory (CPT, [13]). Consequently, RDEUT, either in its own right or as a component of CPT, is one of the most important components of contemporary work to generalize EUT [29, 28].

### Applying the Hierarchical Framework

The elements of the Leslie matrix, *A*_*ij*_, are the determinants of fitness at the intermediate level of Figure 1. We assume that they depend on a valuable but limited resource that can be traded off between fertility and survival across an individual’s life cycle. Stated formally, each Leslie matrix element *A*_*ij*_ is a function of the consumption *x*_*ij*_ allocated to it. *x*_*ij*_ is at the bottom level of Figure 1. We will use a first-order Taylor expansion to assess perturbative changes to the distribution of resources around the baseline optimal distribution **x**_0_. In expressing the Taylor expansion, it is convenient to adopt an indexing notation directly in terms of *F*_*j*_ and *P*_*j*_ rather than the 2D indexing in terms of *A*_*ij*_ (both indexing notations will prove useful below). The fertility in age class *j* is *F*_*j*_ = *A*_1*j*_ whereas the survival in age class *j* is *P*_*j*_ = *A*_*j*+1,*j*_. The corresponding consumption variables are 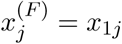 and 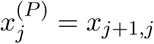. Given this notation, the Taylor expansion for *λ* is

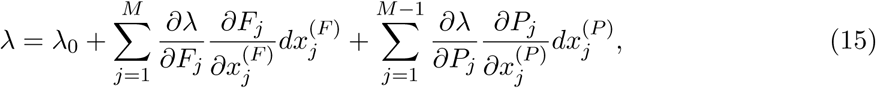

where it is implicit that the partial derivatives are calculated at the baseline distribution of resources **x**_0_ and corresponding baseline Leslie matrix **A**_0_ = **A**(**x**_0_). Caswell [52, 53] derives formulae for the first and second partial derivatives with respect to an element of the Leslie matrix, *∂A*_*ij*_. The first derivative is

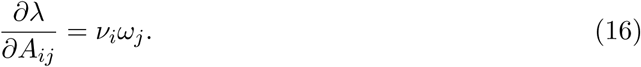

Equation 16 assumes that ***ν*** and ***ω*** have been scaled so that their dot product is 1, ***ν*** ⋅ ***ω*** = 1.

### Pessimism

In this section, we utilize stochastic age-structured life history theory to show that organisms that face uncertainty – arising, e.g., from the interaction of environmental fluctuations interacting with strategic choices – will act as pessimistic decision makers *sensu* RDEUT.^4^ Let *s* index strategies an agent can choose and let *k* index distinct outcomes (for example, achieved survival or fertility in a given time period). The agent faces uncertainty in choosing the strategy since the state of the world is not known when the strategy is chosen, but the agent does know the probability 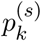 that each outcome will occur given strategy *s*. For the moment, we assume the outcome variable is age-specific fertility in some age class. For outcome *k*, the achieved fertility is *f*_*k*_; we use a lower case *f* to distinguish this fertility from the age-specific fertilities of the Leslie matrix in Equation 3. The mean fertility for strategy *s* is

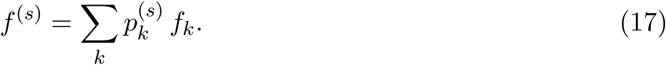

Table 1 illustrates this model for a simple case with three possible strategies and five possible outcomes. What makes these three strategies interesting is that they offer the same expected utility (i.e., fertility) but, as we show next, different evolutionary fitnesses. In particular, strategy *s* = 1 offers the highest fitness because it has the lowest variance and strategy *s* = 3 offers the lowest fitness because it has the highest variance.^5^ Let **A** represent a Leslie matrix in which all elements are fixed except for one stochastic fertility term, 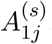, which equals *f*_*k*_ with probability 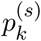 as described above and summarized in Table 1. Tuljapurkar [45] shows that the appropriate fitness measure to use given stochasticity in the matrix elements through time is the long-term logarithmic growth rate. Furthermore, he shows that in a serially-independent environment in which the world state is chosen randomly each time period with no dependence on previous states, the long-term logarithmic growth rate can be approximated by

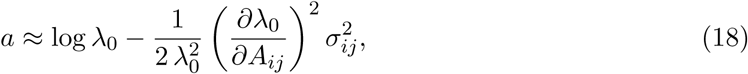

where *λ*_0_ is the growth rate of the (hypothetical) mean phenotype with mean Leslie matrix 〈**A**〉 and 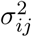 is the variance in the *ij*-th term due to stochastic fluctuations. The crucial insight to be gained from Equation 18 is that the second term is always negative. The effect of stochastic uncertainty, therefore, is to reduce *a* if the mean Leslie matrix is held constant. That is, 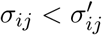 implies 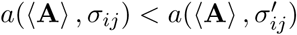, and vice versa. Strategy *s* = 1 in Table 1 is preferred to *s* = 2 and *s* = 2 is preferred to *s* = 3 since variance increases from *s* = 1 to *s* = 3.

**Table 1:**
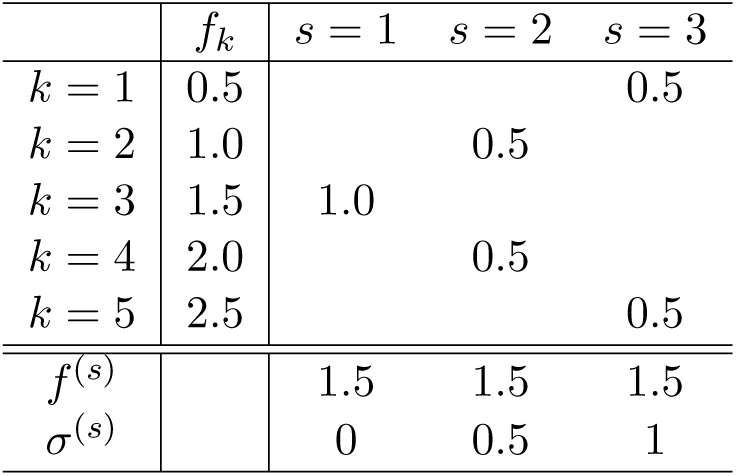
Three strategies with the same expected utility (fertility) but different evolutionary fitnesses. *s* = 1 is preferred to *s* = 2 is preferred to *s* = 3 since variance increases from *s* = 1 to *s* = 3. The table above the double horizontal line summarizes the outcomes for each achieved fertility level, with the first column giving the index *k*, the second column the fertility for each outcome, *f*_*k*_, and the remaining columns the probabilities of each outcome given the chosen strategy *s*. The table below the double horizontal line summarizes, for each strategy *s*, the mean and standard deviation for fertility.

The stage is now set to introduce the economic theory and show that stochastic uncertainty induces deviations from expected utility maximization consistent with pessimistic subjective probability weighting. An expected-utility maximizer will evaluate strategies solely by the mean fertility, *f* ^(*s*)^, which is equivalent to writing *a*^(*s,EUT*)^ = *a*(〈**A**〉, 0), where *a*^(*s,EUT*)^ is the expected utility valuation of the fitness. Symbolically, we can write

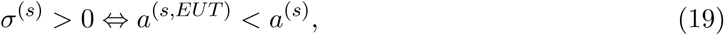

where *⇔* indicates that the lefthand side implies the righthand side and the implication is in both directions. This result is pertinent to RDEUT since the definition of pessimism in RDEUT is that a re-weighted lottery is valued as less than its expected-utility value. For a concave utility function, this is equivalent to assuming that *w*(*p*) ≤ *p* for all *p* [46]. Symbolically, we can write

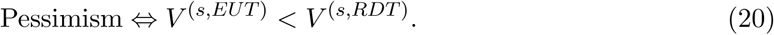

If we assume that natural selection (the principal) has imparted subjective probability weighting to the agent in order to “fix” the optimism of the EUT decision rule, we can posit, by comparing Equation 19 with Equation 20, that natural selection should instill its agents with pessimistic subjective probability weights. It is worth emphasizing that the example in Table 1 is merely for illustration. Equations 18, 19, and 20 apply generally given Equation 10. In the next two sections, we describe and explicate two models to further illustrate the core concepts of our model: a scalar (i.e., one stage) model and a more realistic two stage matrix model. In both models, lotteries that influence fitness are presented to agents, and we show that in both cases pessimistic decision-making maximizes fitness.

### Pessimism: a scalar model

The preceding section utilizes Tuljapurkar’s “small noise approximation” [45] (Equation 10) to establish that serially independent stochastic uncertainty leads to pessimistic probability weighting *sensu* RDEUT.^6^ In this section, we describe a simple model that incorporates only one age class. While this is, of course, unrealistic given the overlapping generations characteristic of the human life history, the simplification nevertheless aids intuition.

The outcome variable is the scalar growth factor, and there are three possible outcomes:

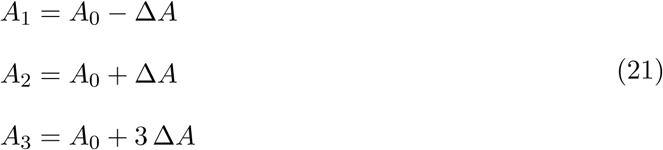

*A*_0_ in Equation 21 is the baseline growth factor, not an additional outcome. There are two strategies, *k* = 1 and *k* = 2. For strategy *s* = 1 outcome *k* = 3, has zero probability and the other two outcomes have probabilities 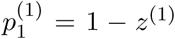 and 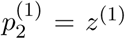, where *z*^(1)^ is a uniform random variable on the interval 0 to 1. For strategy *s* = 2, outcome *k* = 2 has zero probability and the other two outcomes have probabilities 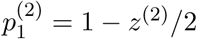 and 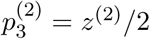, where *z*^(2)^ is (also) a uniform random variable on the interval 0 to 1. Table 2 summarizes these outcomes and probabilities. The agent knows the values of *z*^(1)^ and *z*^(2)^, which are redrawn every time period, and uses probability weighting to choose a strategy each time period. Thus, the agent often (though not always) faces a trade-off between the mean and variance of outcomes induced by the lotteries. The RDEUT values of the collapsed lotteries are (see Table 2 for the contributing lotteries)

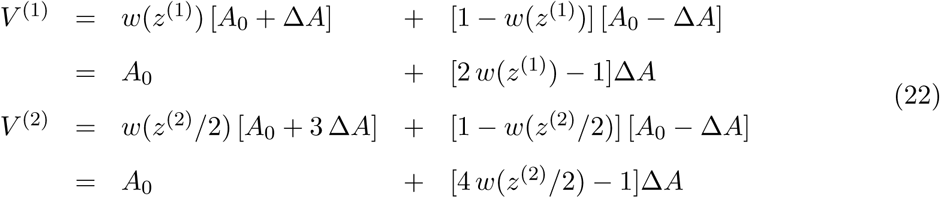

**Table 2:** Payoffs and probabilities for the scalar and matrix model examples. For the matrix model, replace *A*_0_ with *P*_0_ and ∆*A* with ∆*P*. The first column gives the payout index *k*, the second column the payoff for each *k*, and the third and fourth columns the probabilities depending on the strategy *s* that is chosen. *z*^(1)^ and *z*^(2)^ are uniform random variables between 0 and 1 that are drawn anew each time period.

| | Payoff | $s = 1$ | $s = 2$ |
| --- | --- | --- | --- |
| $k = 1$ | $A_0 - \Delta A$ | $1 - z^{(1)}$ | $1 - \frac{z^{(2)}}{2}$ |
| $k = 2$ | $A_0 + \Delta A$ | $z^{(1)}$ | 0 |
| $k = 3$ | $A_0 + 3 \Delta A$ | 0 | $\frac{z^{(2)}}{2}$ |

Setting these equal yields the indifference condition

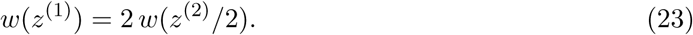

We assume that agents ranks the collapsed lotteries using the function:

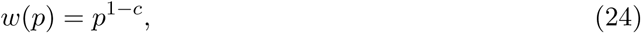

where *c* < 0 implies pessimism, *c* > 0 implies optimism, and *c* < 1. Given this, the indifference curves are lines in the *z*^(1)^-*z*^(2)^ plane,

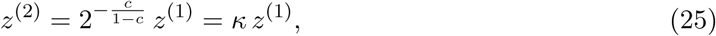

where *κ* = 2^*−c/*(1*−c*)^. Strategy *s* = 2 is preferred to strategy *s* = 1 if *z*^(2)^ is above the indifference curve *g*(*z*^(1)^*, c*). Figure 3 plots these indifference curves for a range of values of *c*.

For this scalar model, the expected log growth rate can be directly calculated by integrating over *z*_1_ and *z*_2_, with strategy *s* = 1 being chosen below the indifference line and strategy *s* = 2 above. For 0 *≤ c <* 1 (or 0 *≤ κ <* 1) the calculation is accomplished by doing the inner integral over *z*^(2)^,

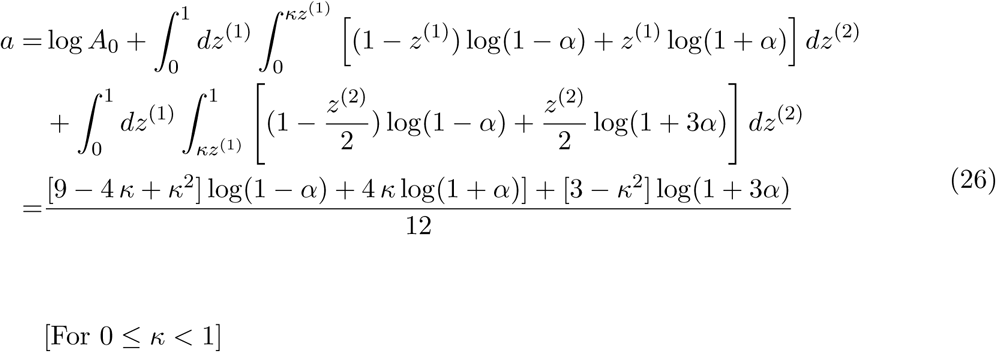

where *α* = ∆*A/A*_0_. For 0 *≤ c* (or 1 *≤ κ ≤* 2) the calculation is accomplished by doing the inner integral over *z*^(1)^,

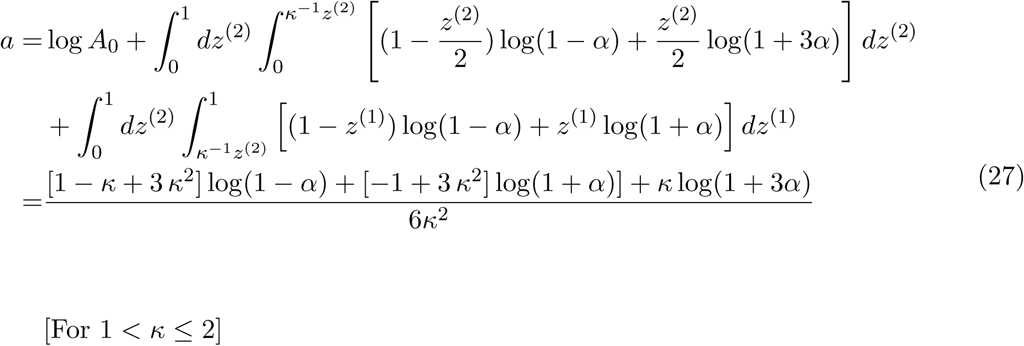

**Figure 3:**
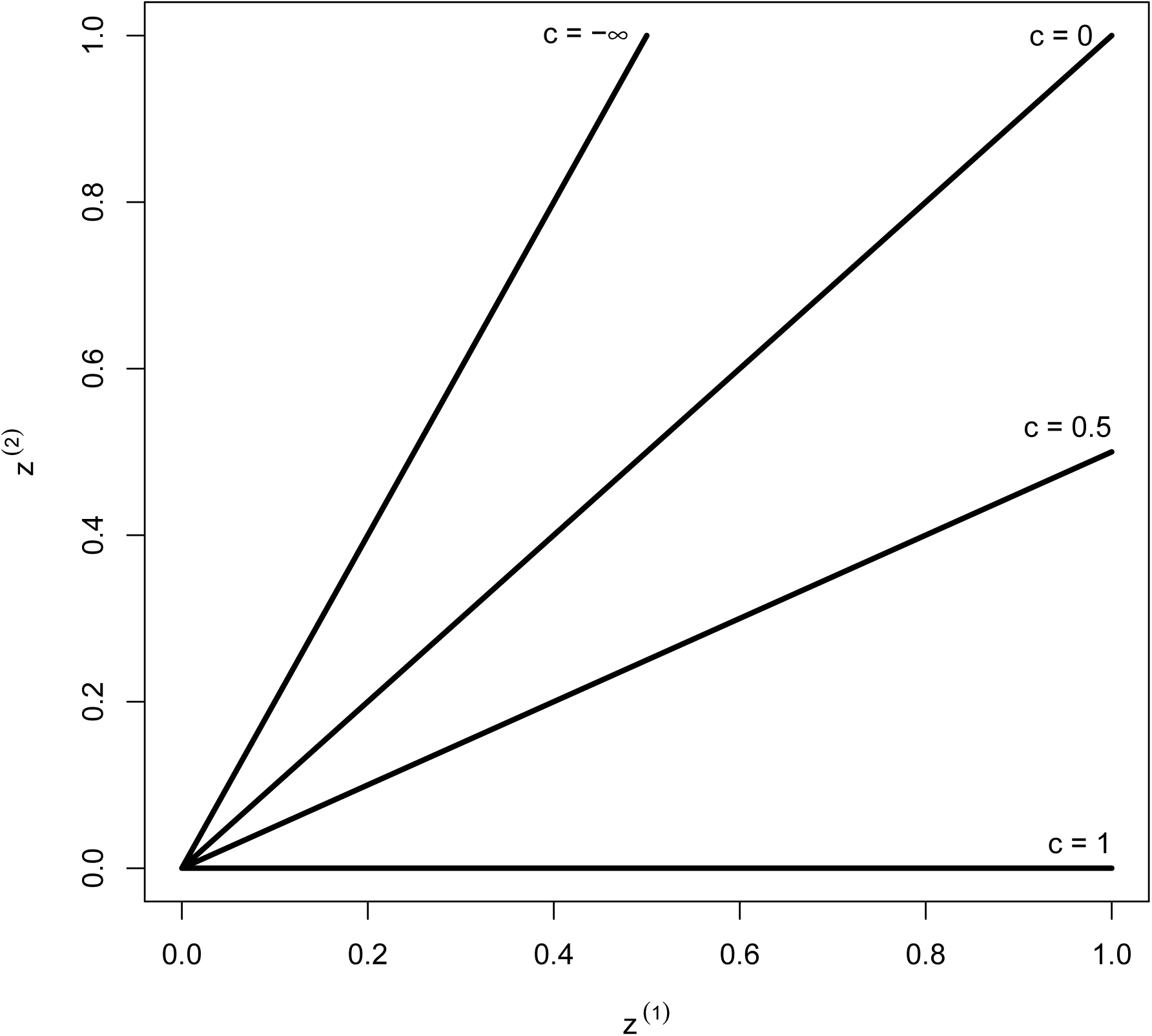
Indifference curves for the scalar model described in the main text. The weighting function is *w*(*p*) = *p*^1*−c*^ (*c* ≤ 1), which yields indifference curves that are lines in the *z*^(1)^-*z*^(2)^ with a slope of 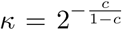 and an intercept of 0. *c* < 0 (*κ* < 1) implies pessimism and *c* > 0 (*κ* > 1) implies optimism. The less risky strategy, *k* = 1, is preferred below the indifference line and the more risk strategy, *k* = 2, is preferred above the indifference line.

The optimal value of *κ* that maximizes *a* is

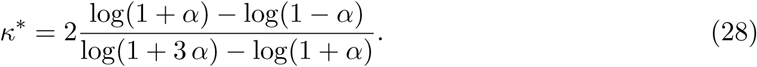

The optimal value of *c* can be found using the relationship

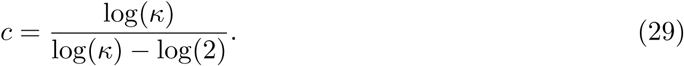

Figure 4 plots *a* as a function of *c* for *A*_0_ = 1.1 and ∆*A* = 0.2; there is an interior maximum at *c* = *−.*27, which indicates that the agent utilizes pessimistic probability weighting (*c* is negative). Figure 5 plots the probability weighting function, *w*(*p*) = *p*^1*−c*^, for this optimal value of *c*.^7^

**Figure 4:**
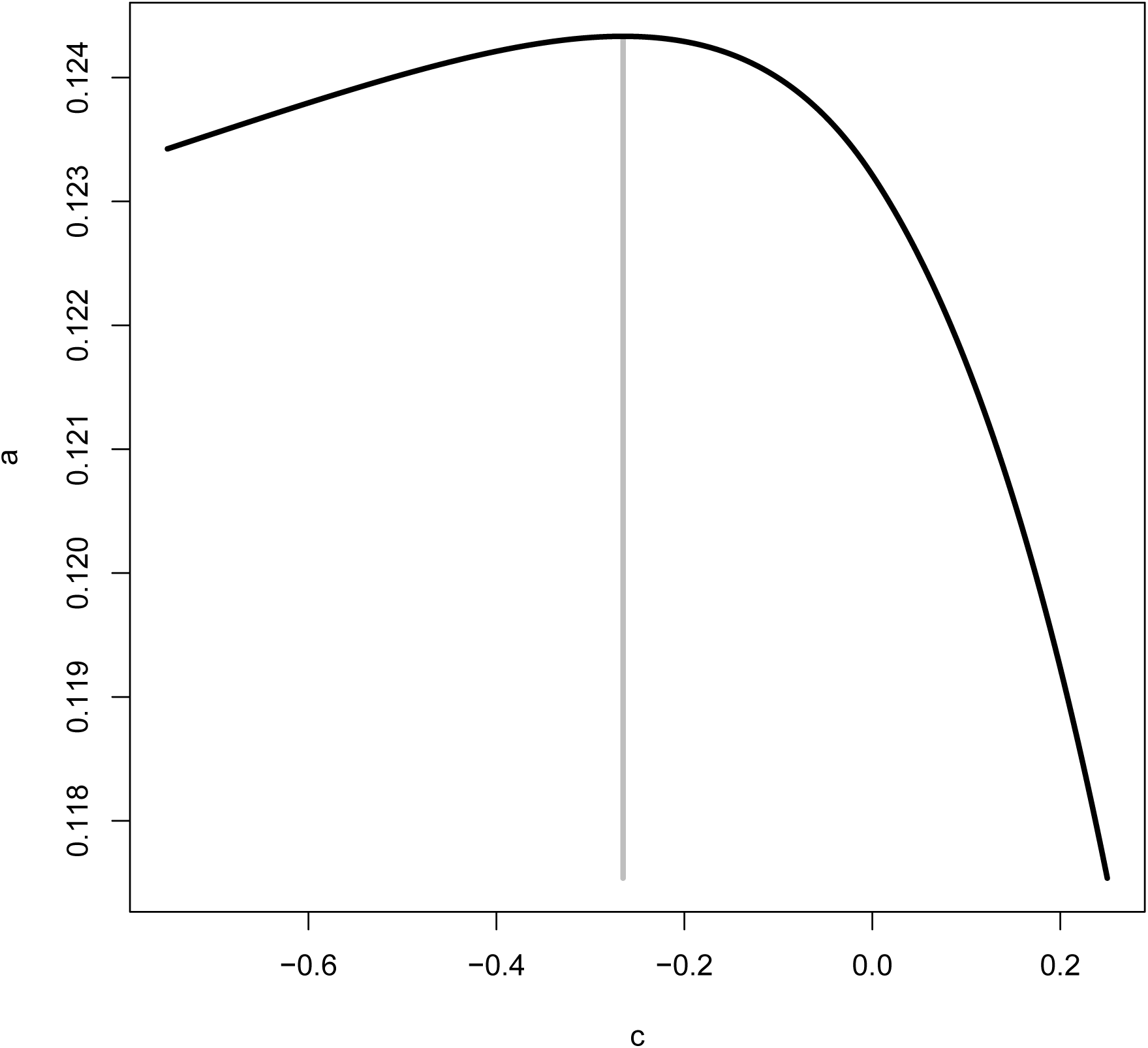
Expected log growth rate (*a*) as a function of the probability weighting parameter *c* for the scalar model. There is an interior maximum at *c* = *−*0.27, marked by the grey vertical line. Figure 5 plots the probability weighting function for this optimal value of *c*. *A*_0_ = 1.1 and ∆*A* = 0.2 for this plot.

**Figure 5:**
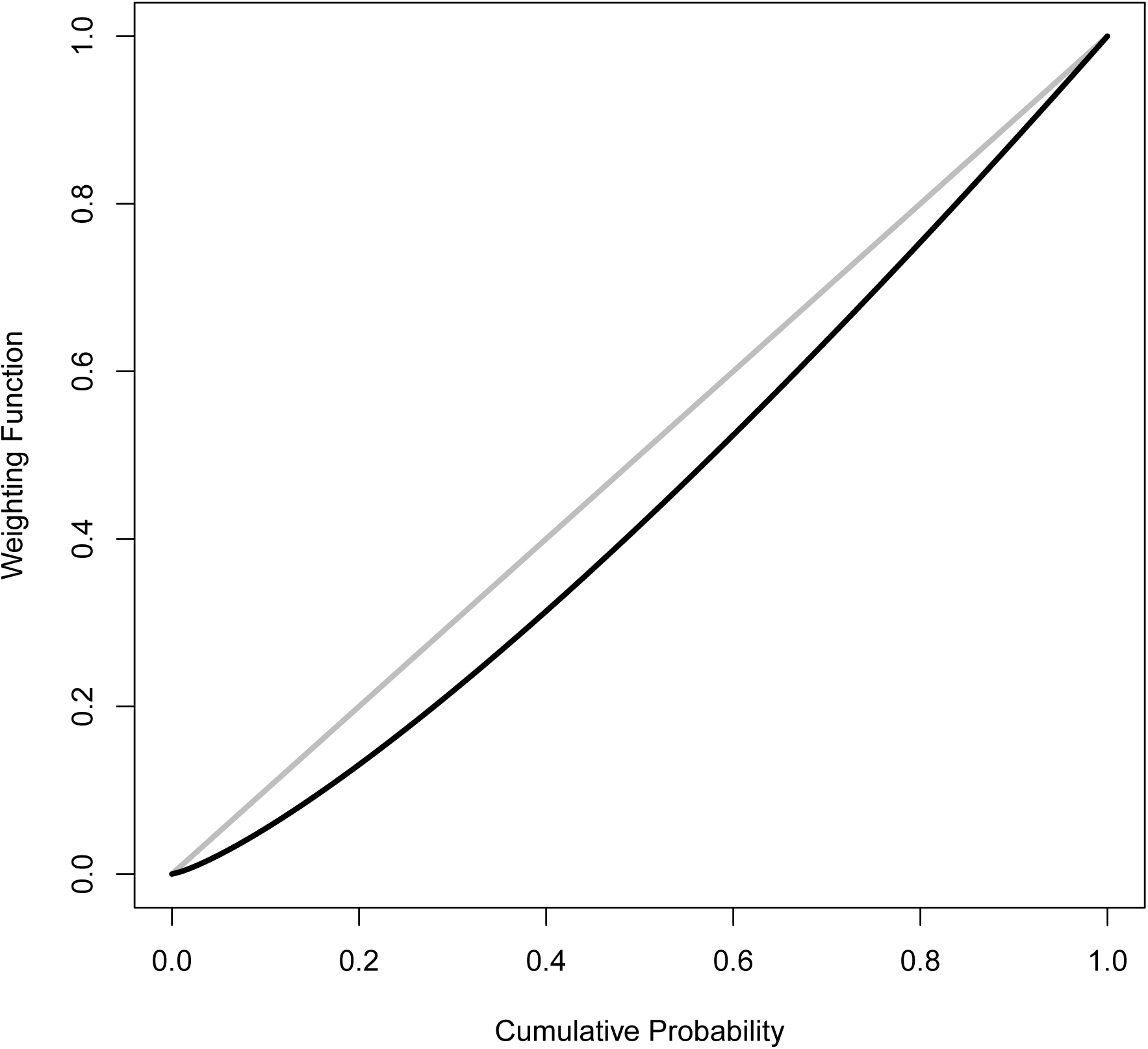
Probability weighting function, *w*(*p*) = *p*^1*−c*^, for the optimal value of *c* = *−*0.27 for the scalar model (see Figure 4). The grey line plots the EUT weighting function *w* = *p*. Since *c* is negative, the agent uses pessimistic probability weighting. *A*_0_ = 1.1 and ∆*A* = 0.2 for this plot.

### Pessimism: a matrix model

In this section, we describe a two stage matrix model for which the population projection matrix is

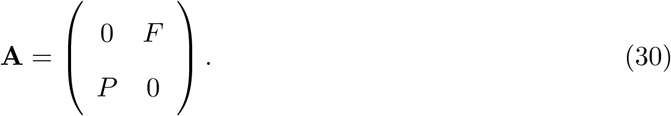

The dominant eigenvalue (i.e., the per-period growth factor) is

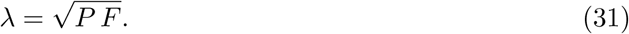

The derivative of *λ* with respect to *P* (used below is)

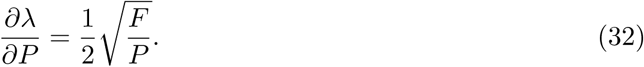

To make this model maximally realistic while still retaining the simplicity of Equation 30, we set the baseline values of *P* and *F* by using the procedure for reduction of the life-ycle graph in [54, Ch. 7], where the reduction is from a full human life cycle with age widths of one year to the two-class form of Equation 30 (the age intervals turn out not to matter). We assume that fertility is zero for the first 15 years, which implies that the reduced values of the juvenile recruitment and growth factor are

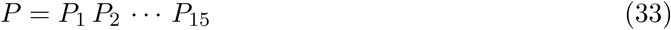

and

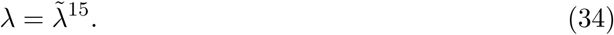

where 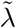 is the baseline per annum population growth factor. Substituting *λ* from Equation 34 in Equation 31 and solving for *F* yields

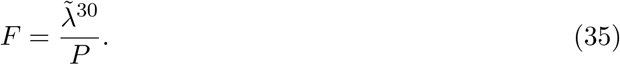

We use a (baseline) juvenile recruitment of *P*_0_ = 0.5 and a per annum growth factor of 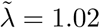 – plausible values for traditional human societies – so that *F* = 1.02^30^/0.5 *≈* 3.6. As above with *A*_0_, *P*_0_ is the baseline payoff, not an additional outcome. We adopt the same payout structure and probabilities as the preceding example (Table 2, with *P*_0_ = 0.5 replacing *A*_0_ and ∆*P* replacing ∆*A*). We adopt ∆*P* = 0.15 for the spread of the gamble.

To apply Equation 10, the mean and variance of *P* must be calculated, which can be done exactly by doing a double integral over *z*^(1)^ and *z*^(2)^. The mean / variance for *κ* ≤ 1 (optimism) and 1 < *κ* ≤ 2 (pessimism) are, respectively,

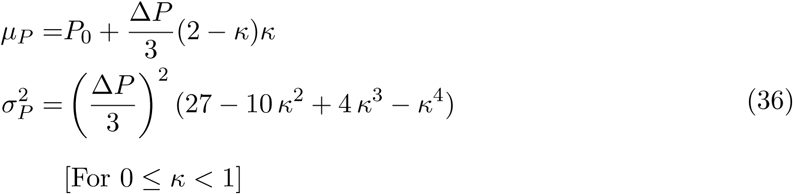

and

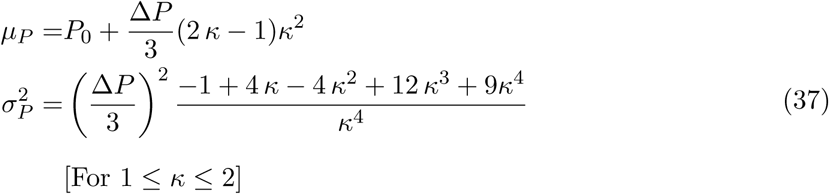

Figure 6 plots *a* as a function of *c*; there is an interior maximum at *c* = *−.*23, which indicates that the agent utilizes pessimistic probability weighting (*c* is negative). Figure 7 plots the probability weighting function, *w*(*p*) = *p*^1*−c*^, for this optimal value of *c*.

**Figure 6:**
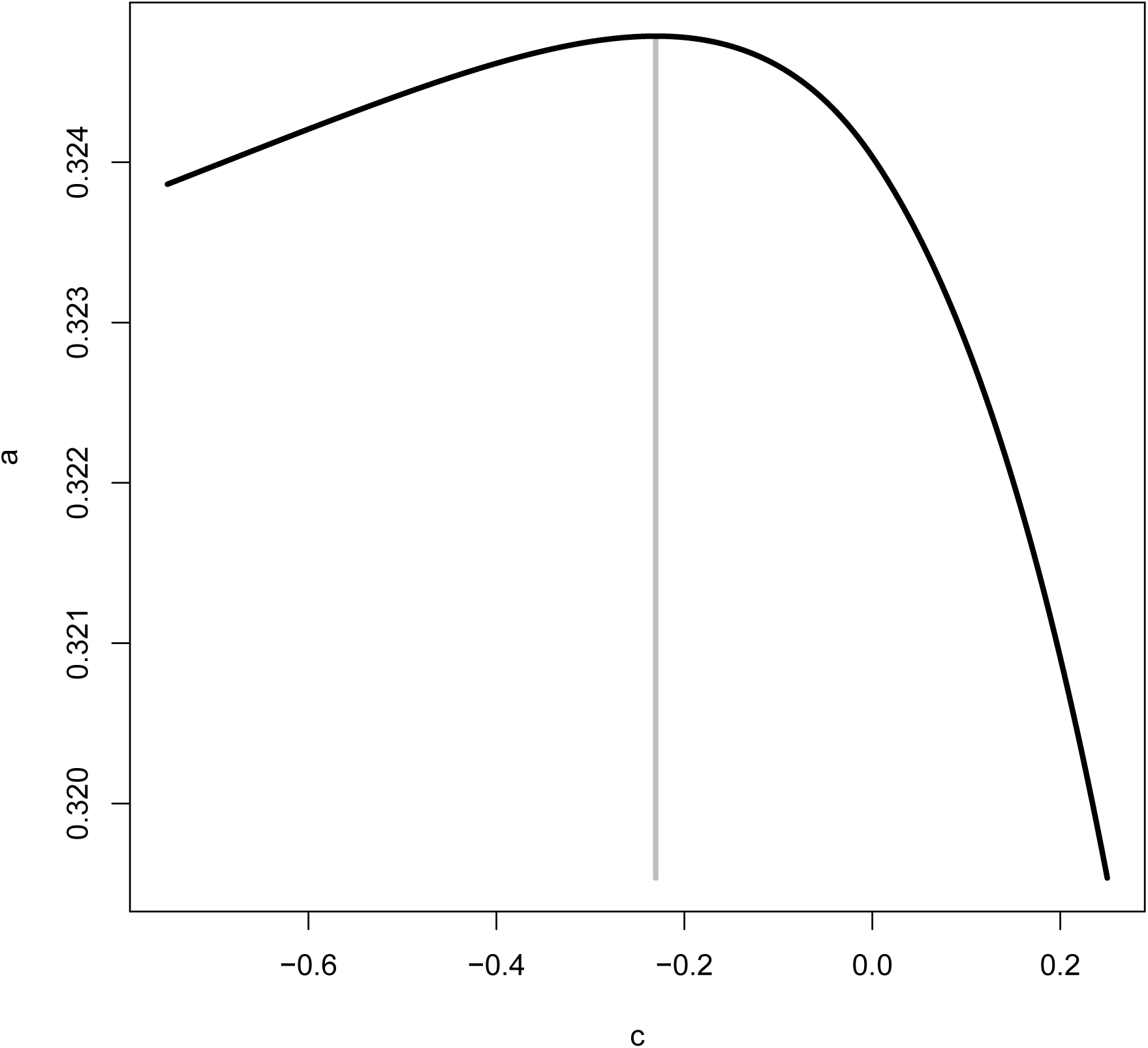
Expected log growth rate (*a*) as a function of the probability weighting parameter *c* for the matrix model. There is an interior maximum at *c* = *−*0.23, marked by the grey vertical line. Figure 7 plots the probability weighting function for this optimal value of *c*. *P*_0_ = 0.5, *F*_0_ = 1.02^30^/0.5 *≈* 3.6, and ∆*P* = 0.15 for this plot.

**Figure 7:**
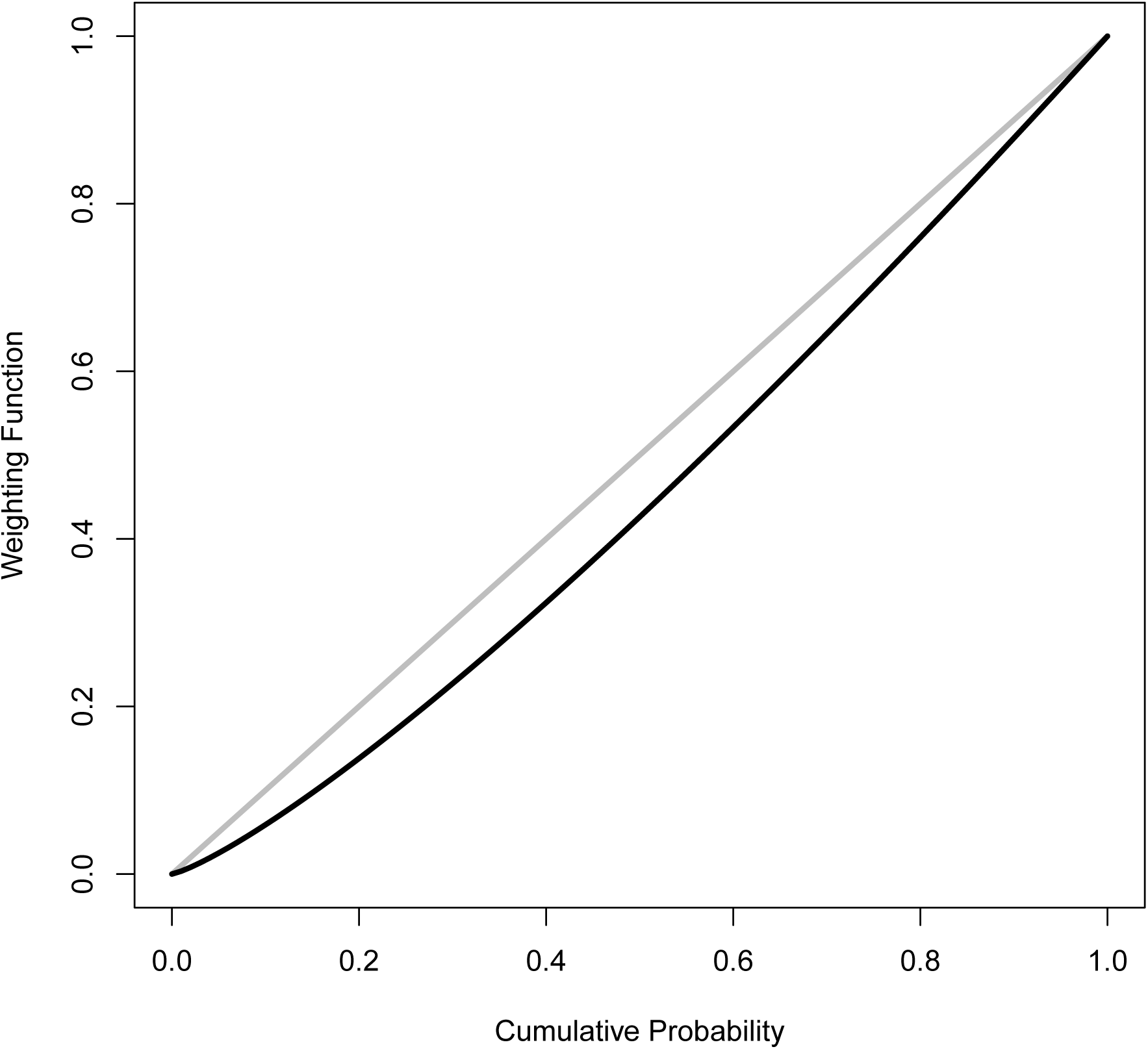
Probability weighting function, *w*(*p*) = *p*^1*−c*^, for the optimal value of *c* = *−*0.23 for the matrix model (see Figure 6). The grey line plots the EUT weighting function *w* = *p*. Since *c* is negative, the agent uses pessimistic probability weighting. *P*_0_ = 0.5, *F*_0_ = 1.02^30^/0.5 *≈* 3.6, and ∆*P* = 0.15 for this plot.

## Discussion

The key scientific finding of this article is the derivation of RDEUT-like pessimistic decision weighting from evolutionary first principles. An explanation for pessimistic probability weighting emerges naturally from coupling utility-maximizing decision-making and fitness maximization. This pessimism arises from a profound intolerance for zeros in key evolutionary parameters which, in turn, arises from a fundamental difference between economic utility and fitness. As we have previously observed, cumulative expected utility is additive across time periods, whereas fitness, and key components of fitness, such as survival, are multiplicative across time, a point made for models of the “survival” of firms by [55]. Consequently, additive decision metrics such as EUT do not necessarily lead to optimal behavior from a fitness standpoint, and can even lead to catastrophic outcomes. This explains the apparent violations of EU optimization observed in subsistence populations such as the herders discussed in the introduction.

There is, in fact, a striking correspondence between the evolutionary intolerance for zeros and the theoretical foundations of RDEUT. John Quiggin, in describing the mentality with which he approached the derivation of RDEUT, writes, “The crucial idea was that the overweighting of small probabilities proposed by Handa and others should be applied only to low probability extreme outcomes, and not to low probability intermediate outcomes” [46, p. 56]. Since low probability extreme outcomes are the key factor driving both the evolutionary intolerance for zeros and the development of RDEU, it may come as no surprise that our evolutionary model predicts the biasing of probability weights for survival and fertility (i.e., utility) in a manner consistent with RDEUT. Aversion to zeros has been used in population biology to explain a range of life-history phenomena which are not favored under standard non-stochastic models such as the regular production of clutches smaller than the most productive clutch [56], delayed reproduction [57], and iteroparity [58].

Our results suggest that evolution will strongly favor pessimistic probability weighting. At first glance, this seems to be at odds with the widely-recognized form of nonlinear weighting associated with CPT, namely, the inverse-S shape indicated in the middle plot of Figure 2. Starting with Tversky and Kahneman [13], weights have been estimated from measures of the certainty-equivalent payoff in experiments [59]. However, these certainty-equivalents conflate risk attitudes resulting from, respectively, the probability weights, *ρ*_*i*_, and the curvature of the utility function, *u*(*x*_*i*_) [60]. While fitness may be a nonlinear function of fertility [34] and fitness in turn depends on underlying consumption variables in a non-linear fashion, the decision maker in our model uses mean fertility as a first-order decision metric, and accounts for the (non-linear) influence of stochasticity via probably weighting. This approach is comparable to the dual model of Yaari [50], though unlike Yaari we do not assume a linear utility function; rather, we account for non-linearity in the utility function as a first order effect, and account for stochasticity via probability weighting.

Our results suggest a new life to long-standing debates on the persistent risk-aversion of agricultural peasants [61, 62, 63] and, more recently, the willingness of the poorest poor to adopt microfinance and other development schemes [64]. The general normative prediction stemming from EUT, and following the foundational paper of Friedman and Savage [65], is that the poorest poor should be willing to take substantial risks to remove themselves from poverty because of the convexity of the putative sigmoid utility function. This logic contributes to the notion that the poorest poor are natural entrepreneurs. However, an increasing body of evidence indicates that the poorest poor are entrepreneurial only to the extent that they lack alternatives such as reliable wage employment. As Banerjee and Duflo [64] write, “are there really a billion barefoot entrepreneurs, as the leaders of MFIs and the socially minded business gurus seem to believe? Or is this just an illusion, stemming from a confusion about what we call an ‘entrepreneur’ ?”

As the decisions of the chronically poor have a far more direct bearing on their survival than, e.g., American undergraduate students in a lab experiment, we expect greater enforcement of the evolutionarily-favored risk preferences. This raises important questions about the ontogeny of risk preference. While models typically treat preferences as fixed, there is experimental evidence that attitudes toward risk are mediated through HPA stress response, both acute [66] and chronic [67, 68], suggesting the possibility for a strong environmental-developmental component to risk preference.

The decision-making of organisms has been shaped by natural selection to render outcomes that are favorable to fitness [69]. While the human brain is considerably more complex than, say, that of a bumblebee, the logic that its capabilities have been shaped by natural selection to execute fitness-enhancing behavior is no less compelling [23]. Biological growth processes are inherently multiplicative, suggesting that decision-making processes shaped by fitness should be especially sensitive to the particulars of persisting when persistence is governed by multiplicative processes. Goats may yield a greater short-term profit, but the mixed herd of goats and camels allows households to persist in the long run [1, 2, 3]. This perspective is supported by evidence that nonhuman decision-makers, for whom the idea that decision-making algorithms have been shaped by selection is more straightforward, are subject to many of the same apparent biases that characterize human decision-making [70]. By demonstrating that fitness maximization in our hierarchical principal-agent model leads directly to pessimistic probability weighting for risky economic decisions linked to fitness – thereby providing a link between evolutionary theory and the rank-dependent family of economic choice models [48, 28] – we hope to stimulate more work on the possible evolutionary foundations of key results from behavioral economics.

## Footnotes

1 Violations of the stationarity of time preferences, another canonical assumption, include the common difference effect and the absolute magnitude effect [14, 15].

2 A virtually identical perspective underlies the so-called “indirect approach,” in which the utility function is defined on proximate goods or outcomes, and the utility function in turn determines the success of an organism [42, 43, 44].

3 [47] may have been the first to suggest the use of subjective probability weights. They showed that when individuals could competitively bid on lotteries with uncertain outcomes they systematically overweighted low probabilities (up to about *p* =. 1) and underweighted high probabilities.

4 Our model generalizes that of [21], who also pointed out that evolution could induce preferences that to do not accord with EUT.

5 The gambles we discuss in this section all depend implicitly on underlying consumption levels that determine fertility. However, the details of the dependence do not impact the results so for simplicity we choose not to explicitly show them.

6 For the scalar model described in this section, a more general result can be established using Jensen’s inequality that does not rely on the small noise approximation, and a similar approach is likely possible for matrix models.

7 Since independent random draws are done for *z*^(1)^ and *z*^(2)^ and the agent can choose the more appealing lottery, the mean log growth rate can exceed log(*A*_0_).

## References

1. Ruth Mace and Alasdair Houston. Pastoralist strategies for survival in unpredictable environments: A model of herd composition that maximises household viability. Agricultural Systems, 31(2):185–204, 1989.

2. Ruth Mace. Pastoralist herd compositions in unpredictable environments: A comparison of model predictions and data from camel-keeping groups. Agricultural Systems, 33(1):1–11, 1990.

3. R. Mace. Nomadic pastoralists adopt subsistence strategies that maximize long-term household survival. Behavioral Ecology and Sociobiology, 33(5):329–334, 1993.

4. Ioannis Karatzas and Steven E Shreve. Methods of Mathematical Finance, volume 39. Springer, 1998.

5. Ole Peters and Murray Gell-Mann. Evaluating gambles using dynamics. Chaos, 26(2), 2016.

6. John von Neumann and Oskar Morgenstern. Theory of Games and Economic Behavior. Princeton University Press, Princeton, NJ, 2nd ed. edition, 1947.

7. L. J. Savage. Foundations of Statistics. Wiley, New York, 1954.

8. Maurice Allais. Le Comportement de l’Homme Rationnel devant le Risque: Critique des Postulats et Axiomes de l’Ecole Americaine. Econometrica, 21:503–546, 1953.

9. Daniel Ellsberg. Risk, ambiguity, and the Savage axioms. The Quarterly Journal of Economics, 75(4):643–669, 1961.

10. Sarah Lichtenstein and Paul Slovic. Reversals of preference between bids and choices in gambling decisions. Journal of Experimental Psychology, 89(1):46–55, 1971.

11. Matthew Rabin. Risk aversion and expected-utility theory: a calibration theorem. Econometrica, 68(5):1281–1292, 2000.

12. Daniel Kahneman and Amos Tversky. Prospect Theory: An analysis of decision under risk. Econometrica, 47(2):263–292, 1979.

13. Amos Tversky and Daniel Kahneman. Advances in Prospect Theory: Cumulative representation of uncertainty. Journal of Risk and Uncertainty, 5:297–323, 1992.

14. George Loewenstein and Drazen Prelec. Anomalies in intertemporal choice: Evidence and an interpretation. The Quarterly Journal of Economics, 107(2):573–597, 1992.

15. Shane Frederick, George Loewenstein, and Ted O’Donoghue. Time discounting and time preference: A critical review. Journal of Economic Literature, 40(2):351–401, 2002.

16. Mark J. Machina. Book review: “rational” decision making versus “rational” decision modelling? Journal of Mathematical Psychology, 24(2):163–175, 1981.

17. Mark J. Machina. Decision-making in the presence of risk. Science, 236(4801):537–543, 1987.

18. Daniel Kahneman. A perspective on judgment and choice: Mapping bounded rationality. American Psychologist, 58(9):697–720, 2003.

19. M. G. Haselton and D. Nettle. The paranoid optimist: An integrative evolutionary model of cognitive biases. Personality and Social Psychology Review, 10(1):47–66, 2006.

20. Rose McDermott, James H. Fowler, and Oleg Smirnov. On the evolutionary origin of prospect theory preferences. The Journal of Politics, 70(02):335–350, 2008.

21. Arthur J. Robson. A biological basis for expected and non-expected utility. Journal of Economic Theory, 68:397–424, 1996.

22. N. Netzer. Evolution of time preferences and attitudes toward risk. American Economic Review, 99(3):937–955, 2009.

23. Leda Cosmides and John Tooby. Better than rational: Evolutionary psychology and the invisible hand. The American Economic Review, 84(2):327–332, 1994.

24. P.M. Todd and G. Gigerenzer. Ecological Rationality: Intelligence in the World. Oxford University Press, Oxford, 2012.

25. Richard C. Lewontin and D. Cohen. On population growth in a randomly varying environment. Proceedings of the National Academy of Sciences of the United States of America, 62(4):1056–1060, 1969.

26. J. H. Gillespie. Natural selection for variances in offspring numbers: A new evolutionary principle. American Naturalist, 111:1010–1014, 1977.

27. Mark J. Machina. Review of ‘generalized expected utility theory: The rank-dependent model’. Journal of Economic Literature, 32(3):1237–1238, 1994.

28. Peter P. Wakker. Prospect Theory: For Risk and Ambiguity. Cambridge University Press, Cambridge, 2010.

29. Chris Starmer. Developments in non-expected utility theory: The hunt for a descriptive theory of choice under risk. Journal of Economic Literature, 38(2):332–382, 2000.

30. Richard Gonzalez and George Wu. On the shape of the probability weighting function. Cognitive Psychology, 38(1):129–166, 1999.

31. James Holland Jones, Rebecca Bliege Bird, and Douglas W. Bird. To kill a kangaroo: understanding the decision to pursue high-risk/high-gain resources. Proceedings of the Royal Society B, 280(1767), 2013.

32. D.W. Norman. Rationalising mixed cropping under indigenous conditions: The example of northern Nigeria. Journal of Development Studies, 11:3751, 1974.

33. J.N. Hobcraft, J. MacDonald, and S. Rutstein. Child spacing effects on infant and early child mortality. Population Index, 49(4):585–618, 1983.

34. James Holland Jones and Rebecca Bliege Bird. The marginal valuation of fertility. Evolution and Human Behavior, 35(1):65–71, 2014.

35. Joghn P. Shelton. The Cost of Renting versus Owning a Home. Land Economics, 44(1):59–72, 1968.

36. Larry Samuelson and Jeroen Swinkels. Information, evolution and utility. Theoretical Economics, 1:119–142, 2006.

37. A. R. Rogers. Evolution of time preference by natural selection. American Economic Review, 84(3):460–481, 1994.

38. Ken G. Binmore. Game Theory and the Social Contract, volume 1. MIT Press, Cambridge, MA, 1994.

39. Arthur J. Robson and Larry Samuelson. The Evolutionary Foundations of Preferences. In Jess Benhabib, Alberto Bisin, and Matthew O. Jackson, editors. Handbook of Social Economics, volume 1A, pages 221–310. Elsevier, 2011.

40. P. W. Glimcher. Proximate mechanisms of individiaul decision-making behavior. In D. S. Wilson and A. Kirman, editors. Complexity and Evolution: Toward a New Synthesis for Economics, pages 85–96. MIT Press, Cambridge, 2016.

41. George C. Williams. Adaptation and natural selection. Princeton University Press, Princeton, 1966.

42. Werner Güth and Menahem Yaari. Explaining recicrocal behavior in simple strategic games: An evolutionary approach. In U. Witt, editor, Explaining Process and Change: Approaches to Evolutionary Economics, pages 23–34. University of Michigan Press, Ann Arbor, 1992.

43. Werner Güth. An evolutionary approach to explaining cooperative behavior by reciprocal incentives. International Journal of Game Theory, 24(4):323–344, 1995.

44. Eddie Dekel, Jeffrey C. Ely, and Okan Yilankaya. Evolution of preferences. The Review of Economic Studies, 74(3):685–704, 2007.

45. Shripad Tuljapurkar. Population dynamics in variable environments. III. evolutionary dynamics of r-selection. Theoretical Population Biology, 21:141–165, 1982.

46. John Quiggin. Generalized Expected Utility Theory: The Rank-Dependent Model. Kluwer, Norwell, MS, 1993.

47. Malcolm G. Preson and Philip Baratta. An experimental study of the auction-value of an uncertain outcome. American Journal of Psychology, 61(2):183–193, 1948.

48. John Quiggin. A theory of anticipated utility. Journal of Economic Behavior and Organization, 3:323–343, 1982.

49. David Schmeidler. Subjective probability and expected utility without additivity. Econometrica, 57:571–587, 1989.

50. Menahem E. Yaari. The dual theory of choice under risk. Econometrica, 55:95–115, 1987.

51. Uzi Segal. Some remarks on Quiggin’s anticipated utility. Journal of Economic Behavior & Organization, 8(1):145–154, 1987.

52. H. Caswell. A general formula for the sensitivity of population growth rate to changes in life history parameters. Theoretical Population Biology, 14:215–230, 1978.

53. Hal Caswell. Second derivatives of population growth rate: Calculation and applications. Ecology, 77(3):870–879, 1996.

54. H. Caswell. Matrix Population Models: Construction, Analysis and Interpretation. Sinauer, Sunderland, MA, 2nd edition, 2001.

55. Roy Radner. Economic survival. Econometric Society Monographs, 29:183–209, 1998.

56. Mark S. Boyce and C. M. Perrins. Optimizing great tit clutch size in a fluctuating environment. Ecology, 68(1):142–153, 1987.

57. S. Tuljapurkar. Delayed reproduction and fitness in variable environments. Proceedings of the National Academy of Sciences, USA, 87(3):1139–1143, 1990.

58. S.H. Orzack and S. Tuljapurkar. Population dynamics in variable environments. VII. demography and evolution of iteroparity. American Naturalist, 133(6):901–923, 1989.

59. Nathaniel T. Wilcox. Random expected utility and certainty equivalents: mimicry of probability weighting functions. Journal of the Economic Science Association, 3(2):161–173, 2017.

60. Enrico Diecidue and Peter P. Wakker. On the intuition of rank-dependent utility. Journal of Risk and Uncertainty, 23(3):281–298, 2001.

61. M. Lipton. Theory of optimising peasant. Journal of Development Studies, 4(3):327–351, 1968.

62. S.L. Popkin. The Rational Peasant: The Political Economy of Rural Society in Vietnam. University of California Press, Berkeley, 1979.

63. J. Henrich and R McElreath. Are peasants risk-averse decision makers? Current Anthropology, 43(1):172–181, 2002.

64. A.V. Banerjee and E. Duflo. Poor Economics: A Radical Rethinking of the Way to Fight Global Poverty. PublicAffairs, New York, 2011.

65. M. Friedman and L. J. Savage. The Utility Analysis of Choices Involving Risk. The Journal of Political Economy, 56(4):279–304, 1948.

66. J. Cahlikova and L. Cingl. Risk preferences under acute stress. Experimental Economics, 20(1):209–236, 2017.

67. Narayanan Kandasamy, Ben Hardy, Lionel Page, Markus Schaffner, Johann Graggaber, Andrew S. Powlson, Paul C. Fletcher, Mark Gurnell, and John Coates. Cortisol shifts financial risk preferences. Proceedings of the National Academy of Sciences, 111(9):3608–3613, 2014.

68. P. Kusev, H. Purser, R. Heilman, A. J. Cooke, P. Van Schaik, V. Baranova, R. Martin, and P. Ayton. Understanding risky behavior: The influence of cognitive, emotional and hormonal factors on decision-making under risk. Frontiers in Psychology, 8, 2017.

69. L.A. Real. Animal choice behavior and the evolution of cognitive architecture. Science, 253(5023):980–986, 1991.

70. Laurie R. Santos and Alexandra G. Rosati. The evolutionary roots of human decision making. Annual Review of Psychology, 66(1):321–347, 2015.

